# TMEM180 contributes to colorectal cancer proliferation through intracellular metabolic pathways

**DOI:** 10.1101/2020.07.16.207712

**Authors:** Takahiro Anzai, Shinji Saijou, Yoshitsugu Ohnuki, Hiroshi Kurosawa, Masahiro Yasunaga, Yasuhiro Matsumura

## Abstract

TMEM180, a novel colon cancer–specific membrane protein with a 12-transmembrane topology, is upregulated at low oxygen. Previously, we established a humanized monoclonal antibody against TMEM180 aimed at clinical trials. Prior to such trials, it is necessary to clarify the function of this molecule in cancer. To compare SW480 human colon cancer cells and their TMEM180-knockdown derivatives, we analyzed proliferation and oxygen consumption, and also performed phosphorylation proteomics, metabolomics, and next-generation sequencing. The results revealed that TMEM180 promoted the growth of colon cancer but had almost no effect on oxygen consumption or expression of phosphorylated proteins. By contrast, glycolysis differed dramatically between SW480 and TMEM180-knockdown cells. TMEM180 promotes nitric oxide synthesis, suggesting that it promotes glucose metabolism and glutamine metabolism, thereby contributing to cancer growth. Overall, the results of this study support the clinical development of an anti-TMEM180 antibody.

## Introduction

Colorectal cancer (CRC) is the third leading cause of cancer death and has a high incidence and mortality worldwide(1). Consequently, there is considerable incentive to identify new target molecules for the diagnosis and treatment of CRC. We previously identified a new membrane protein, TMEM180, that is highly expressed in CRC, and successfully developed the anti-TMEM180 monoclonal antibody (mAb) for future clinical use(2). We reported that TMEM180 is upregulated under low-oxygen conditions and may play an important role in the uptake or metabolism of glutamine and arginine in cancer cell proliferation(2). We also showed that *Tmem180*-knockout mice do not exhibit embryonic, neonatal, or postnatal lethality(2). Recently, we found that TMEM180 has 12 transmembrane domains and that its N- and C-termini are exposed extracellularly(3). TMEM180 was inferred to be a cation symporter(3), but its biological function in CRC cells remains unclear.

In this study, we sought to clarify the function of TMEM180 by analysis of oxygen consumption, phosphorylated protein proteomics, next-generation sequencing, and metabolomics.

## Materials and Methods

### Cells and cell cultures

SW480 cells were purchased from American Type Culture Collection. Cells were cultured in DMEM low-glucose medium (Wako) supplemented with 10% FBS (Thermo Fisher Scientific) and 1% penicillin–streptomycin–amphotericin B suspension (Wako) at 37°C under a 5% CO_2_ atmosphere. Clones of SW480 cells harboring stable knockdown of *TMEM180* were established as described previously(2). Lentiviral transduction particles were used to generate stable knockdown cells (Sigma-Aldrich, MISSION TRC clones TRCN0000243137 for *TMEM180* knockdown and SHC005V for *eGFP* knockdown; the latter was used as the Mock control).

### Quantitative real-time RT-PCR

To measure the level of *TMEM180* mRNA in stable knockdown cells, quantitative real-time RT-PCR was performed as described previously(2). The relative expression of *TMEM180* was normalized against expression of *GAPDH*.

### Cell proliferation assay

For the CCK-8 assay, SW480 cells were seeded at 250 cells per well in 200 μL culture medium in 96-well cell culture plates (Corning) on day 0. After culturing for 2, 4, 6, or 7 days at 37°C, 5% CO_2,_ cell proliferation was measured using the Cell-Counting Kit 8 (Dojindo, CCK-8). CCK-8 solution was added (10 μL per well), and the plate was incubated for 3 h. Absorbance of each well was measured at 450 nm. For the anchor-independent cell proliferation assay, SW480 cells were seeded at 125 cells per well in 200 μL culture medium in round-bottom Ultra Low Attachment (ULA) 96-well plates (Corning). The ULA plates were centrifuged at 100 × g for 3 min, and the cells were cultured at 37°C under 5% CO_2_ with shaking at 80 rpm. Images were acquired on a Keyence Microscope BZX-700 (Keyence) on day 0 (12 h), day 2, day 4, day 6, and day7 after cells were seeded.

### Measurement of respiratory rate

A respiratory rate measurement system using a spinner flask was constructed as described in Fig.2A. The spinner flask was filled with 60 mL of DMEM low-glucose medium and incubated at 35°C under a 4% CO_2_ atmosphere with gentle stirring at 80 rpm. Cell suspension (5 mL of 7–8×10^7^ cells) was inoculated, and dissolved oxygen concentration in the spinner flask was measured with an oxygen probe (DKK Co., DOL-10). First, the volumetric oxygen transfer coefficient *k_L_a* was determined as follows. Dissolved oxygen in the medium was purged by nitrogen substitution. After the dissolved oxygen concentration (*C*) was lowered to near zero, the medium was re-oxygenated using the gas atmosphere in the incubator, described above. Dissolved oxygen concentration was recorded as a function of time. The slope of the curve, the derivative 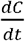, corresponds to the oxygen transfer rate, and is expressed by equation (a).

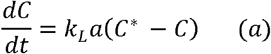

*C\**: saturated dissolved oxygen concentration [mg-O_2_·L^−1^]

*C*: dissolved oxygen concentration [mg-O_2_·L^−1^]

*k_L_a*: oxygen transfer capacity coefficient [min^−1^]

*t*: time [min]

Integration of equation (a) under the initial conditions *t* = 0 and *C* = *C_0_* yields equation (b).

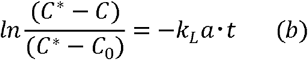

Based on equation (b), *k_L_a* was calculated to be 0.064 [min^−1^] from the slope of a straight line on a semi-log plot. For the cell culture system, a term for respiration rate (*r_o2_*) is added to the equation (a).

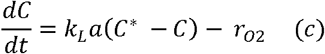

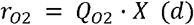

*r_o2_*: respiration rate [mg-O_2_·L^−1^·min^−1^]

*Q_O2_*: specific respiration rate [mg-O_2_·cells^−1^·min^−1^]

*X*: cell concentration in the measurement system [cells·L^−1^]

At steady state, the oxygen supply rate (*k*_*L*_*a* (*C*∗ − *C*)) and the respiration rate (*r_o2_*) are balanced, and the dissolved oxygen concentration (*C*) is constant, i.e., 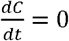. Therefore, equation (e) can be obtained from equation (c).

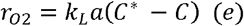

Respiratory rate (*r_o2_*) was calculated by substituting the value of *C* at steady state into equation (e). *r_o2_* divided by cell concentration is the specific respiratory rate (*Q_O2_*).

### Preparation of phosphoprotein-enriched extracts

Cultured cells were washed three times with PBS, and pellets were harvested in sample buffer containing 4% SDS, 125 mM Tris-HCl (pH 6.8), and 0.04% bromophenol blue. Protein solutions were boiled 95°C for 5 min and stored at −20°C. Samples were dissolved in a sample lysis solution containing 7 M urea, 2 M thiourea, 4% (w/v) 3-[(3-cholamidopropyl)dimethyammonio]-1-propanesulfonate (CHAPS), 1% (w/v) dithiothreitol (DTT), 2% (v/v) Pharmalyte, and 1 mM benzamidine, and homogenized using a PowerGen125 motor-driven homogenizer (Thermo Fisher Scientific). Total proteins were extracted for 15 min at room temperature with vortexing, and the extract was centrifuged at 12,000 × g for 20 min at 25°C. The PhosPro Phosphoprotein enrichment kit (Genomine) was used to isolate phosphoproteins from total protein extracts. Subsequently, the sample was mixed with 750 μL of delipidation solution (methanol:chloroform = 600:150), vortexed vigorously for 5 min, and centrifuged at 15,000 × g for 10 min to promote phase separation. The middle phase containing the protein disk was recovered, and the upper and lower phases were discarded. The protein disk was washed twice in ~1 mL methanol. The protein pellet was completely air-dried or dried in an oven and dissolved in a buffer for 2D electrophoresis.

### 2D electrophoresis

IPG strips (Genomine) were reswelled for 12–16 h at room temperature in a solution containing 7 M urea, 2 M thiourea, 2% CHAPS, 1% DTT, and 1% Pharmalyte. Isoelectric focusing was performed using 800 μg protein sample per strip on a MultiPhor II system (Amersham Biosciences) at 20°C. The voltage was sequentially increased from 0.15 to 3.5 kV over 3 h to allow entry of the sample, followed by maintenance at 3.5 kV over 9 h, with focusing completed after 96 kV-h. IPG strips were incubated for 10 min in equilibration buffer (50 mM Tris-HCl, pH 6.8 containing 6 M urea, 2% SDS, and 30% glycerol), first with 1% DTT and second with 2.5% iodoacetamide. Equilibrated strips were loaded onto SDS-PAGE gels (10 cm × 12 cm, 10–16%), and SDS-PAGE was performed on a Hoefer DALT 2D system (Amersham Biosciences) at 20°C for 1.7 kV-h. Gels were fixed with a solution containing 40% (v/v) ethanol and 10% (v/v) acetic acid for 1 h, and then stirred three times for 30 min in a rehydration solution (5% (v/v) ethanol and 5% (v/v) acetic acid in distilled water. Phosphoproteins were visualized using ProQ Diamond phosphoprotein gel stain (Invitrogen) for 2 h, and then washed with ProQ Diamond phosphoprotein destaining solution (Invitrogen) for 60 min. Gel images were acquired using DIVERSITY (Syngene). The gels were washed with distilled water three times and stained with Coomassie Brilliant Blue G-250 (Invitrogen). Images were acquired on a Duoscan T1200 (Agfa). Intensities of individual protein spots were normalized against the total intensities of all valid spots. Analysis was performed using the PDQuest 2D analysis software (Bio-Rad). Protein spots that exhibited a significant (≥2-fold) change in expression between WT and KD were selected for further analysis.

### PMF analysis

Selected protein spots were excised from the gel and enzymatically digested in-gel essentially as previously described(4) using trypsin (Promega). Gel pieces were washed with 50% acetonitrile to remove SDS, salt, and stain; dried to remove solvent; and then rehydrated with trypsin and incubated for 12 h at 37°C. The proteolytic reaction was terminated by addition of 5 μL 0.5% trifluoroacetic acid. Tryptic peptides were recovered by combining the aqueous phase from several extractions of the gel pieces with 50% aqueous acetonitrile. After concentration, the peptide mixture was desalted using C18ZipTips (Millipore), and peptides were eluted in 1–5 μL of acetonitrile. An aliquot of this solution was mixed with an equal volume of saturated solution of α-cyano-4-hydroxycinnamic acid in 50% aqueous acetonitrile / 0.1% TFA, and 1 μL of the mixture was spotted onto a target plate. Proteins were subjected to MALDI-TOF analysis (Bruker Daltonics, Microflex LRF 20) as described previously(5). The search program MASCOT, developed by Matrix Science, was used for protein identification by peptide mass fingerprinting.

### Metabolite extraction

Cells were grown to a density of 1–5 × 10^6^ cells per 10-cm dish, washed three times with PBS, and cultured in DMEM low-glucose medium without FBS. After 3 h, culture medium was removed from the dish, and the cells were washed twice in 5% mannitol solution, first with 10 mL and next with 2 mL. Metabolites were extracted from 2.7–3.0 × 10^6^ cells with 800 μL methanol and 550 μL Milli-Q water containing Internal Standard Solution (Human Metabolome Technologies [HMT]) for CE-MS, and with 1300 μL ethanol containing Internal Standard Solution for LC-MS. Extracts for CE-MS were transferred into a microfuge tube and centrifuged at 2,300 × g at 4°C for 5 min. To remove proteins, the extracts were centrifugally filtered through a 5-kDa cutoff filter (Millipore) at 9,100 × g at 4°C for 2 h. Extracts were stored at −80°C until analysis. Before measurement, extracts for CE-MS were centrifugally concentrated and resuspended in 50 μL of Milli-Q water for measurement. Extracts for LC-MS were mixed with 1,000 μL Milli-Q water and sonicated for 5 min while cooling on ice, and the supernatant was collected by centrifugation (4,400 × g, 4°C, 5 min). It was dissolved in 200 μL of 50% aqueous isopropanol solution (v/v) and used for the measurement.

### Metabolomic analysis

Targeted quantitative analysis of metabolites was performed by HMT using capillary electrophoresis time-of-flight mass spectrometry (CE-TOF/MS), capillary electrophoresis triple quadrupole mass spectrometry (CE-QqQMS), and liquid chromatography time-of-flight mass spectrometry (LC-TOF/MS). Analytic methods were described previously(6). CE-TOFMS measurement was performed on an Agilent CE-TOFMS system, CE-MS/MS measurement was performed on an Agilent CE system with Agilent 6460 TripleQuad LC/MS, and LC-TOFMS measurement was performed using an Agilent 1200 series RRLC system SL with Agilent LC/MSD TOF (all machines: Agilent Technologies). Peaks detected by CE-TOF/MS and LC-TOF/MS were extracted using MasterHands ver2.17.1.11 (Keio University), and peaks detected by CE-MS/MS were extracted using MassHunter Quantitative Analysis B.06.00 (Agilent Technologies). Migration time (for CE-MS), retention time (for LC-MS), *m/z*, and peak area were obtained from the software. The peaks were annotated according to the HMT metabolite database based on their *m/z* values with the Migration times or retention times. The obtained relative area value was converted into an absolute quantitative value using a standard substance. The peak area value corrected by the internal standard substance was used for quantitative conversion. A calibration curve consisting of three points was created for each metabolite, and the concentration was calculated. Hierarchical cluster analysis (HCA) and principal component analysis (PCA) were performed by HMT using in-house analysis software developed by the company.

### RNA-seq and enrichment analysis

Total RNA was extracted using the RNeasy Mini kit (Qiagen). RNA quality checking, library preparation, and sequencing on the HiSeq 4000 platform (Illumina) were performed by Eurofins Genomics. All samples had an RNA Integrity Number (RIN)◻>◻9.7. Trimmomatic(7) ver0.36 was used to remove adaptor sequences and low-quality reads from the sequencing data. BWA(8) ver0.7.17 was used to map reads onto the human reference genome assembly GRCh38.p12. Count files produced by featureCounts(9) were normalized and statistically analyzed by the edgeR package using TCC-GUI(10). Differentially expressed genes (DEGs) compared with each group were identified with a q value < 0.1. Enrichment analysis was performed on upregulated or downregulated DEGs using Metascape(11).

## Statistical analysis

Significant differences between two groups were determined using Student’s t-test for Fig. 1A, B, D, Fig. 5D, Fig. S3C, and Fig. S4C, and Welch’s t-test for Fig. 4C.

**Figure 1.**
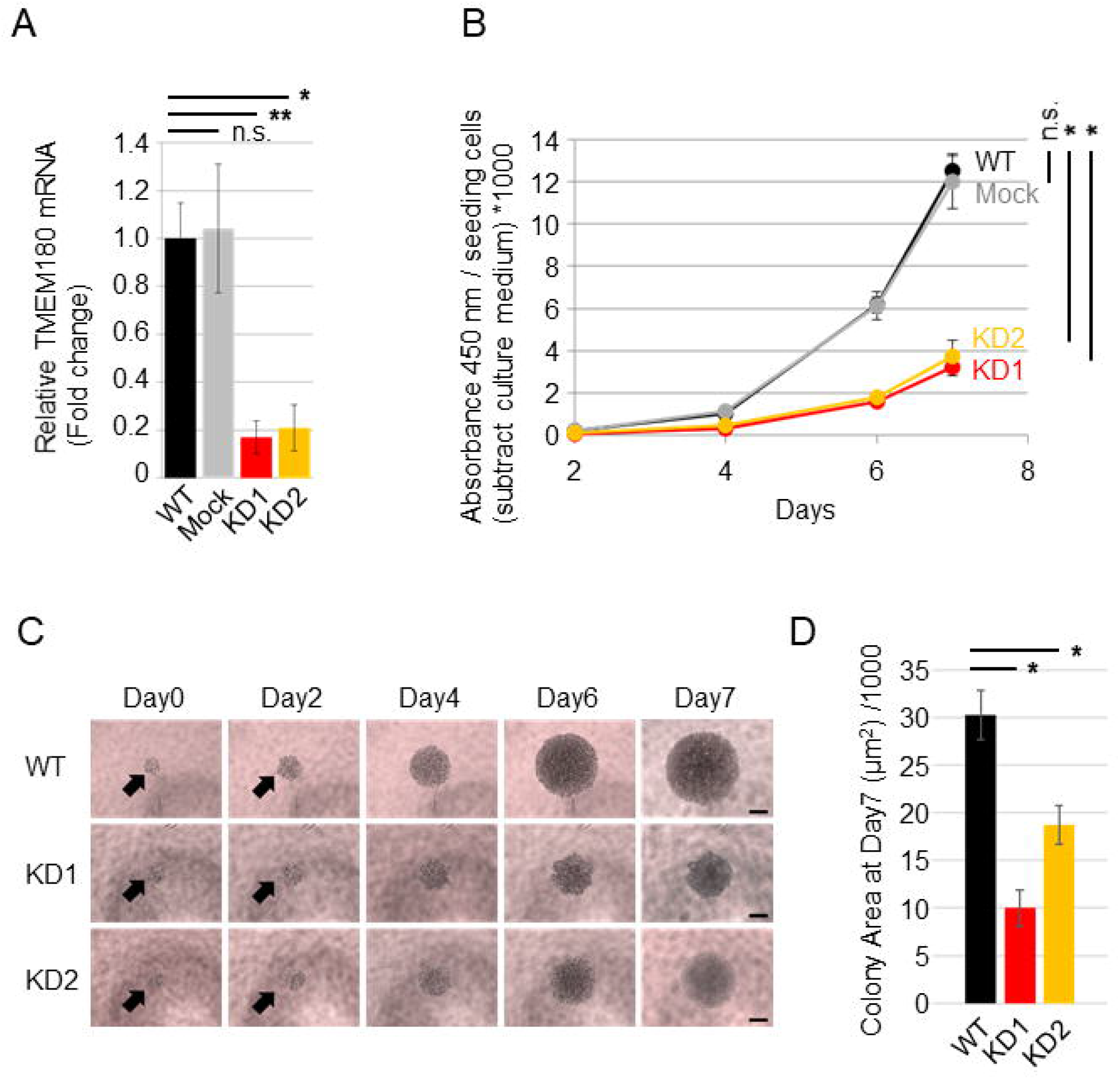
TMEM180 knockdown suppresses SW480 cell proliferation. A. Relative expression of TMEM180 in SW480 (WT), SW480 expressing control shRNA (Mock), and SW480 expressing TMEM180-specific shRNA (KD1 and KD2) was measured by qRT-PCR. *P < 0.05, **P < 0.01, n.s. = not significant. Bars = SD. B. Effect of TMEM180 knockdown on cell proliferation. CCK-8 assays indicated that TMEM180 knockdown suppressed cell proliferation. Absorbance at 450 nm was multiplied by 1000 after subtraction of background value (culture medium without cells), and then normalized against cell number. Statistical analysis was performed using the value from Day 7. *P < 0.05, n.s. = not significant. Bars = SD. C. Typical microscope images of SW480 WT, KD1, and KD2 cells taken on Day 0 (12 h after seeding cells), Day 2, Day 4, Day 6, and Day7. Scale bars: 200 μm. D. Cell colony area on Day 7, calculated using ImageJ1.52 from six independent wells. **P < 0.01, Bars = SD.

## Results

### TMEM180 gene knockdown suppress cell proliferation of SW480 colon cancer cells

To explore molecular function of TMEM180 in cancer cells, we established TMEM180-knockdown SW480 cell clones using shRNA (Fig. 1A). To evaluate effects of TMEM180 gene knockdown, we performed cell proliferation assay of SW480 cells. Cell proliferation was significantly lower in TMEM180-knockdown cell clones (KD1 and KD2) than in the parental cell line (WT) or cells expressing a control shRNA (Mock) (Fig. 1B). Next, we compared anchoring independent cell proliferation between WT and KD cells. Anchorage-independent cell proliferation was also significantly suppressed in TMEM180-knockdown cell clones (KD1 and KD2) relative to WT (Fig. 1C, D).

### TMEM180 gene knockdown does not contribute to SW480 cell respiration rate

Oxygen concentration profiles in the spinner flasks inoculated with SW480 WT and SW480 KD cells are shown in Fig. 2A. Specific respiration rate (Q_O2_), oxygen consumption rate per unit cell, calculated from the equilibrium value of dissolved oxygen concentration (C). There is no significant difference in Q_O2_ between SW480 WT and SW480 KD cells (Fig. 2B). The results suggest that TMEM180 gene knockdown does not have an effect on cell respiration. It is supposed that TMEM180 is not involved in the function of oxygen uptake in cells or mitochondria. The correlation between cell proliferation and oxygen consumption is well known in cancer cells(12). In this study, it was found that cell proliferation was promoted in SW480 WT cells. Moreover, SW480 WT cells formed spherical cell aggregates that were larger in size than SW480 KD cells. The increase in cell number or in size of cell aggregate will increase the overall oxygen consumption, but it may not be directly linked to the oxygen consumption rate per unit cell (Q_O2_). Formation of three-dimensional cell aggregate will generate low-oxygen environment in the central region of them. The expression of TMEM180 gene under low-oxygen environment may be advantageous to cell proliferation to form larger cell aggregates such as solid tumors. These observations suggest that TMEM180 is not involved in oxygen uptake in cells or mitochondria.

**Figure. 2.**
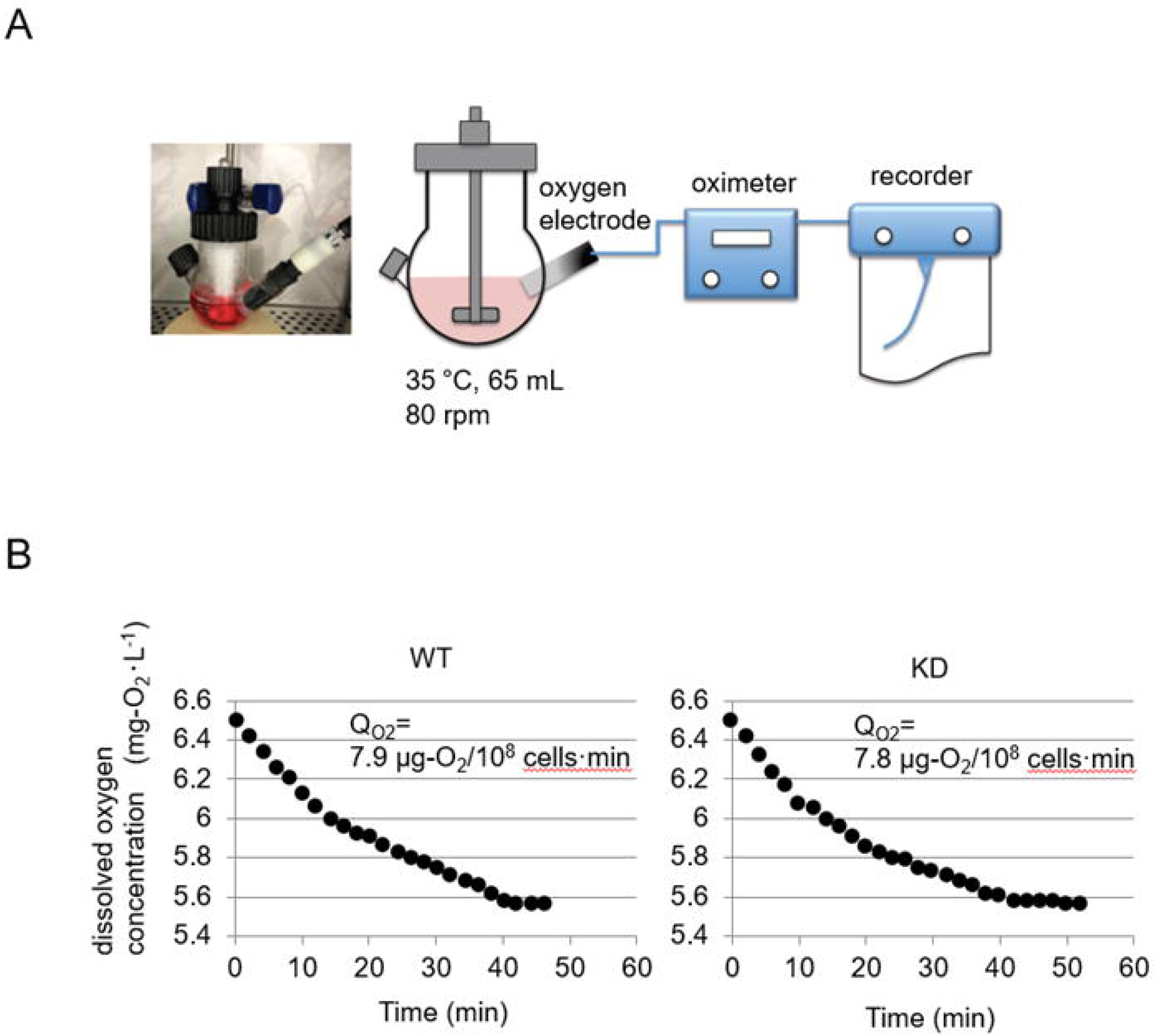
Respiration rate of SW480 cells. A. Oxygen concentration profiles in spinner flasks inoculated with cells. B. Cell respiration data plot and respiration rate of WT (left) and KD (right) cells.

### TMEM180 gene knockdown has little effect on phosphoprotein expression

To investigate changes in phosphorylation proteins associated with TMEM180 gene knockdown, we performed phospho-proteomics analysis between WT and KD cells. We identified enriched phosphoproteins by 2D gel electrophoresis followed by staining with CBB and ProQ Diamond (Fig. 3A). Only two protein spots differed in intensity between WT and KD cells (Fig. 3A,B). By searching the database using the Mascot server, we identified spot 1 as epididymis luminal protein 176 and spot 2 as alpha-enolase (Fig. 3B). Based on the UniProtKB(13) annotation, epididymis luminal protein 176 (UniProtKB: V9HVZ7) has an unknown function, but may belong to the actin family. The other protein, alpha-enolase (ENO1) (UniProtKB: P06733), is a key enzyme in the glycolytic pathway(14), and its expression is correlated with cancer progression or metastasis(15,16). It is not known whether phosphorylation of ENO1 is related to cancer. These results indicate that TMEM180 has a minimal effect on phosphoprotein expression.

**Figure. 3.**
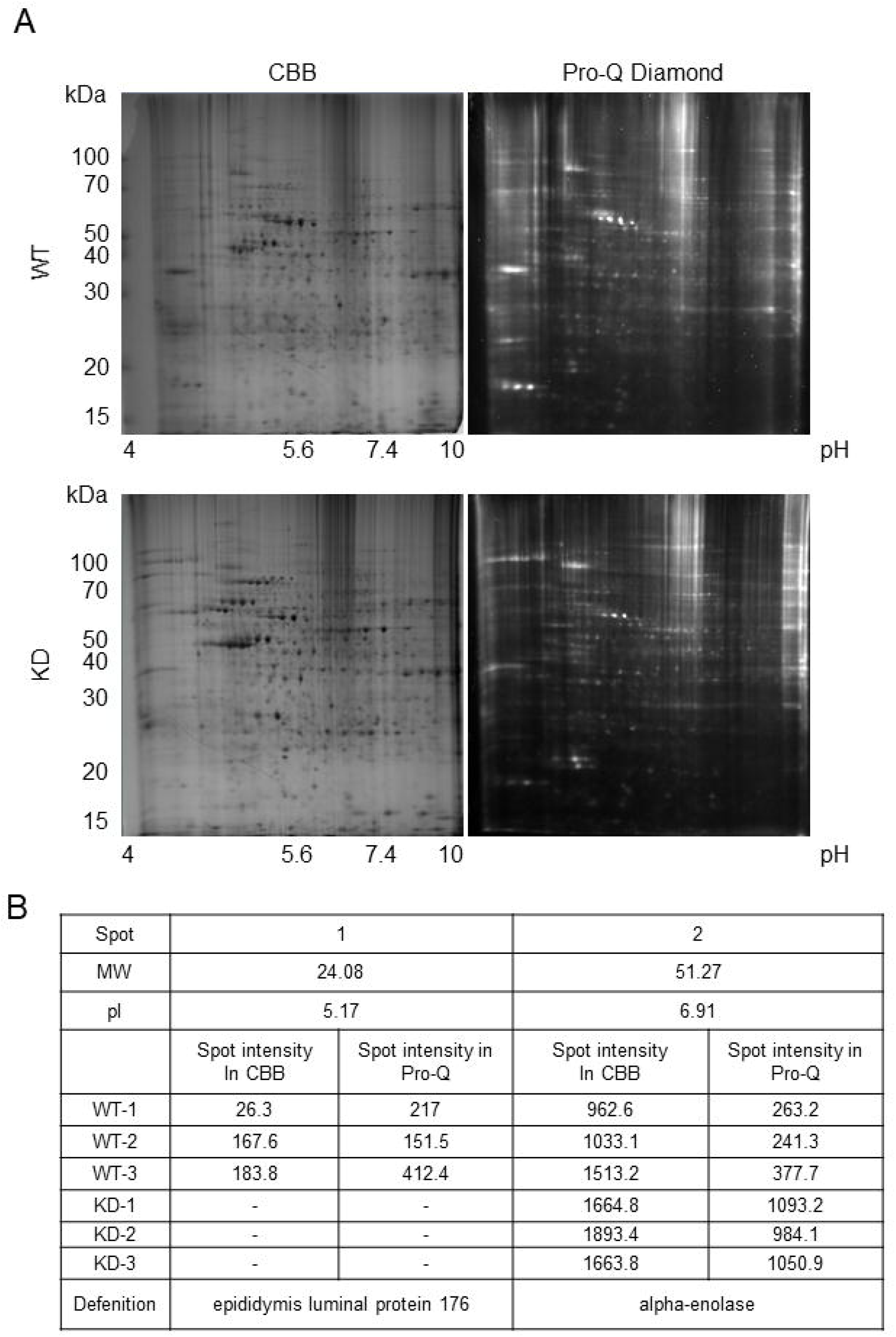
Phosphoprotein patterns of SW480 cells, monitored by 2D gel electrophoresis. A. 2D gel images from one of three independent experiments are shown. Upper panels represent WT cells, and lower panels represent KD cells. Left panels show Coomassie Brilliant Blue (CBB) staining, and right panels show ProQ Diamond staining. Y-axes indicate apparent molecular mass (kDa), and X-axes indicate pH. B. Results of differential spot analysis. MW is the predicted molecular weight of the spot, and pI is the predicted isoelectric point of the spot.

**Figure 4.**
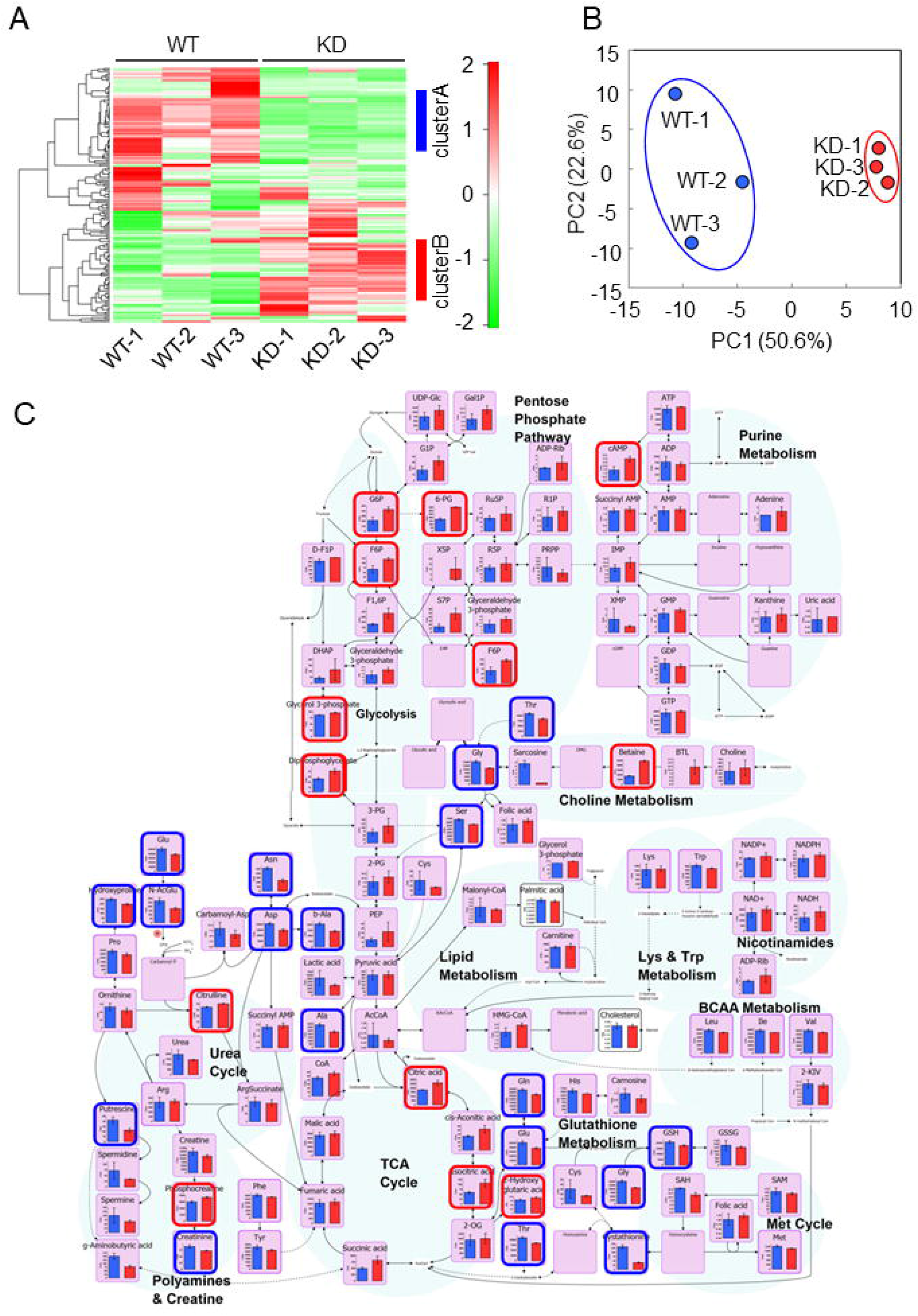
Metabolomics analysis of SW480 cells. A. Heat map representation of metabolome profiles analyzed by hierarchical clustering analysis. Cluster A (blue line) and cluster B (red line) are shown on the right edge of the figure. B. Score plot of principal component 1 (PC1) versus principal component 2 (PC2) from the principal component analysis (PCA). Clustering of WT samples (blue circle) and KD samples (red circle) is shown. C. Graphical representation of metabolites mapped to known pathways of glycolysis, lipid, and amino acid metabolism. Bar graphs represent the amounts of metabolites (exceptions: palmitic acid and cholesterol are relative values) in WT (blue) and KD (red) cell samples. Metabolites that were significantly more abundant in WT are highlighted with blue squares, and those that were more abundant in KD are highlighted with red squares.

**Figure 5.**
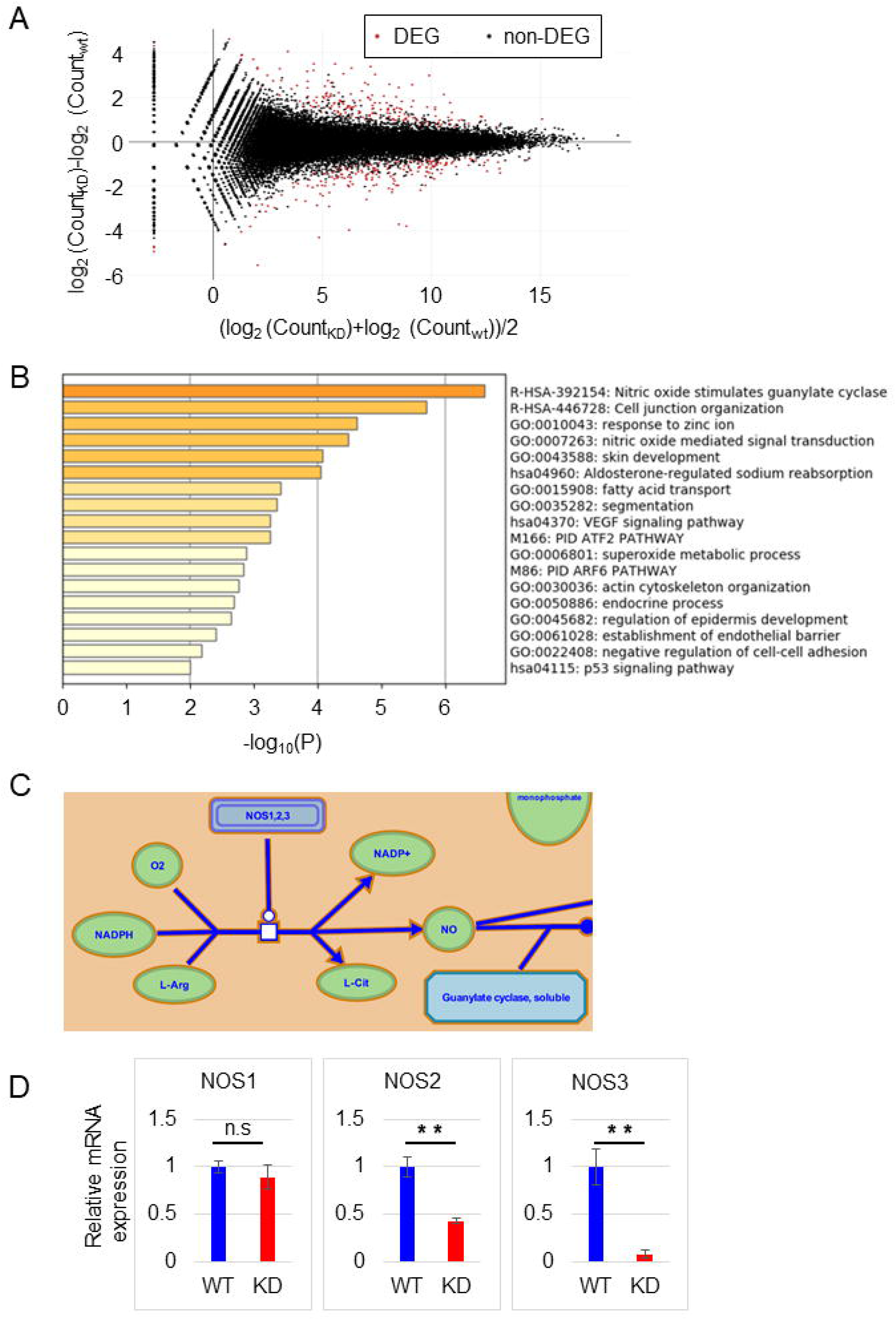
RNA-seq analysis of SW480 cells. A. MA plot comparing WT and KD cells. Genes colored in red were selected as DEGs using an FDR q-value threshold of 0.1. B. Function enrichment analysis of DEG correlated with TMEM180 KD from between WT and KD cell samples. C. Nitric oxide (NO) stimulates guanylate cyclase (Reactome R-HSA-392154). NO generation, the first step of the pathway, is shown. D. Relative expression of *NOS1, NOS2,* and *NOS3* from our RNA-seq data. WT value is defined as 1. **P < 0.01, n.s. = not significant. Bars = SD.

### TMEM180 gene knockdown cause variation in metabolites around glycolysis

To investigate the metabolite profile changes associated with TMEM180 gene knockdown, we performed capillary electrophoresis time of flight mass spectrometry (CE-TOF/MS), capillary electrophoresis-triple quadrupole mass spectrometry (CE-QqQMS) and liquid chromatograph time of flight mass spectrometry (LC-TOF/MS). In this analysis, we detected a total of 161 metabolites (CE-MS: 44 and 57 in cation and anion mode, respectively; LC-MS: 46 and 14 metabolites in positive and negative mode, respectively). The heat map in Fig. 4A represents two major clusters. In cluster A, several amino acids and acylcarnitine-related metabolites were present at higher levels in WT cells than in KD cells. In cluster B, levels of phospholipid- and glycolysis-related metabolites were elevated in KD cells relative to WT cells (Supplementary Table S1). The score plot of the principal component analysis (PCA) revealed a clear separation between WT and KD along the first principal component (PC1) axis (Fig. 4B). Glycolytic metabolites and lysophospholipids were included among the top 30 metabolites with positive values of the principal component score (Supplementary Table S2), and amino acids and acylcarnitine were included in the top 30 metabolites with negative values (Supplementary Table S3). The detected metabolites were mapped to known pathways of glycolysis and amino acid metabolism (Fig. 4C). Related metabolites upstream of glycolysis were more abundant in KD cells (Fig. 4C, highlighted in red square). However, there was no difference in the amount of pyruvic acid or lactate. In addition, levels of amino acids such as Asn, Asp, Gln, Glu, and Ser that flow into the glycolysis pathway were significantly higher in WT cells (Fig. 4C, highlighted in blue square). Based on these results, knockdown of TMEM180 appeared to affect glycolysis and amino acid metabolism.

### TMEM180 gene knockdown cause downregulation of nitric oxide synthase and glutaminase

In RNA-seq analysis, the MA plot revealed that 165 genes were downregulated, and 160 genes were upregulated, with q-value < 1.0 (Fig. 5A). In biological enrichment analysis of the downregulated genes using Metascape(11), nitric oxide (NO) stimulation of guanylate cyclase (Reactome(17) Gene Sets) was top-ranked (Fig. 5B). In this pathway, nitric oxide synthase (NOS) produces NO, which oxidizes a guanidine nitrogen of L-arginine (Fig. 5C and Fig. S1). NO is an essential molecule involved in several pathophysiological processes in mammals(18). Three isoforms of NOS have been identified: neuronal (nNOS or NOS1), inducible (iNOS or NOS2), and endothelial (eNOS or NOS3)(18). We found that expression of *NOS2* and *NOS3* was higher in WT cells than in KD cells, whereas expression of *NOS1* did not differ (Fig. 5D). On the other hand, enrichment analysis of upregulated genes revealed functions related to developmental or morphological processes (Fig. S2).

We next analyzed public TCGA data (https://www.cancer.gov/tcga) to search for genes that correlate with TMEM180 expression. Correlation analysis between normal and primary tumors was performed using UALCAN(19). GLS2 was the only relevant metabolic enzyme out of the top 10 positively correlated genes (Fig.S3A). The correlation between TMEM180 (x-axis) and GLS2 (y-axis) is visualized as Fig. S3B. We confirmed that expression of GLS2 was higher in WT cells than in KD cells (Fig. S3C). We previously reported that TMEM180-knockdown cells could not grow in serum-free medium without glutamine and arginine(2). Thus, TMEM180 may play an important role in the uptake or metabolism of glutamine and arginine during tumor growth and proliferation(2). Based on these findings, we hypothesized that TMEM180 is related to both NO-related metabolism and glutamine metabolism.

## Discussion

Our findings indicated that TMEM180 is involved in the growth of CRC cells (Fig. 1), as confirmed in KD cells derived from other CRC cell lines (data not shown). To investigate the mechanisms involved in proliferation, we compared respiratory rate, phosphorylation signals, and metabolites between SW480 WT and its TMEM180-knockdown derivative. The results revealed that TMEM180 does not contribute to cellular respiration rate (Fig. 2) and has little effect on phosphoprotein expression (Fig. 3). On the other hand, TMEM180 caused variation in metabolites related to glycolysis (Fig. 4). RNA-seq analysis revealed that NOS and glutaminase genes were downregulated in TMEM180 KD cells (Fig. 5 and Fig. S3). NO may influence glucose and glutamine utilization in tumor cells directly or through the activation of oncogenic pathways(18). Glutaminase catalyzes the conversion of glutamine into glutamate. There are two subtypes of glutaminase: GLS (kidney-type) and GLS2 (liver-type)(20). Elevated expression of GLS has been observed in several types of cancer(21). In CRC, high expression of GLS is correlated with poor prognosis(22). By contrast, overexpression of GLS2 acts in an anti-oncogenic manner in liver and brain cancer(23),(24). Hence, we analyzed TCGA data from CRC using GEPIA2(25). The results revealed that expression of neither GLS nor GLS2 was associated with poor prognosis (Fig. S4A,B). In our RNA-seq data, expression of GLS was unchanged (Fig. S4C). Thus, the relationship between expression of GLS2 and cancer proliferation in patients remains controversial.

In TMEM180 KD cells, the upstream of glycolytic pathway is enhanced, but there was no change in the amount of pyruvic acid, and the amount of lactate was higher in WT cells. Activation of glycolysis by NO, activation of the glutamine pathway by high expression of GLS2, and amino acid uptake were observed in SW480 WT cells, suggesting that proliferation was promoted in SW480 WT cells with high TMEM180 expression (Fig. 6). Recently, Mei et al. reported that siRNA-mediated knockdown of MFSD13A/TMEM180 promotes proliferation of pancreatic cancer cell lines(26). On the other hand, we showed that shRNA-mediated knockdown of TMEM180 suppressed proliferation of SW480 cells (Fig. 1). Thus, the role of TMEM180 in proliferation may differ depending on the type of cancer. Further studies in different types of cancer are needed to resolve this potential contradiction.

In conclusion, we showed that TMEM180 contributes to the growth of CRC by altering metabolism, rather than signal transduction or mitochondrial function. In future work, it will be important to identify the substrate that is transported by coupling with cations. We anticipate that these findings will help elucidate more precisely the role of TMEM180 in cancer cells.

**Figure 6.**
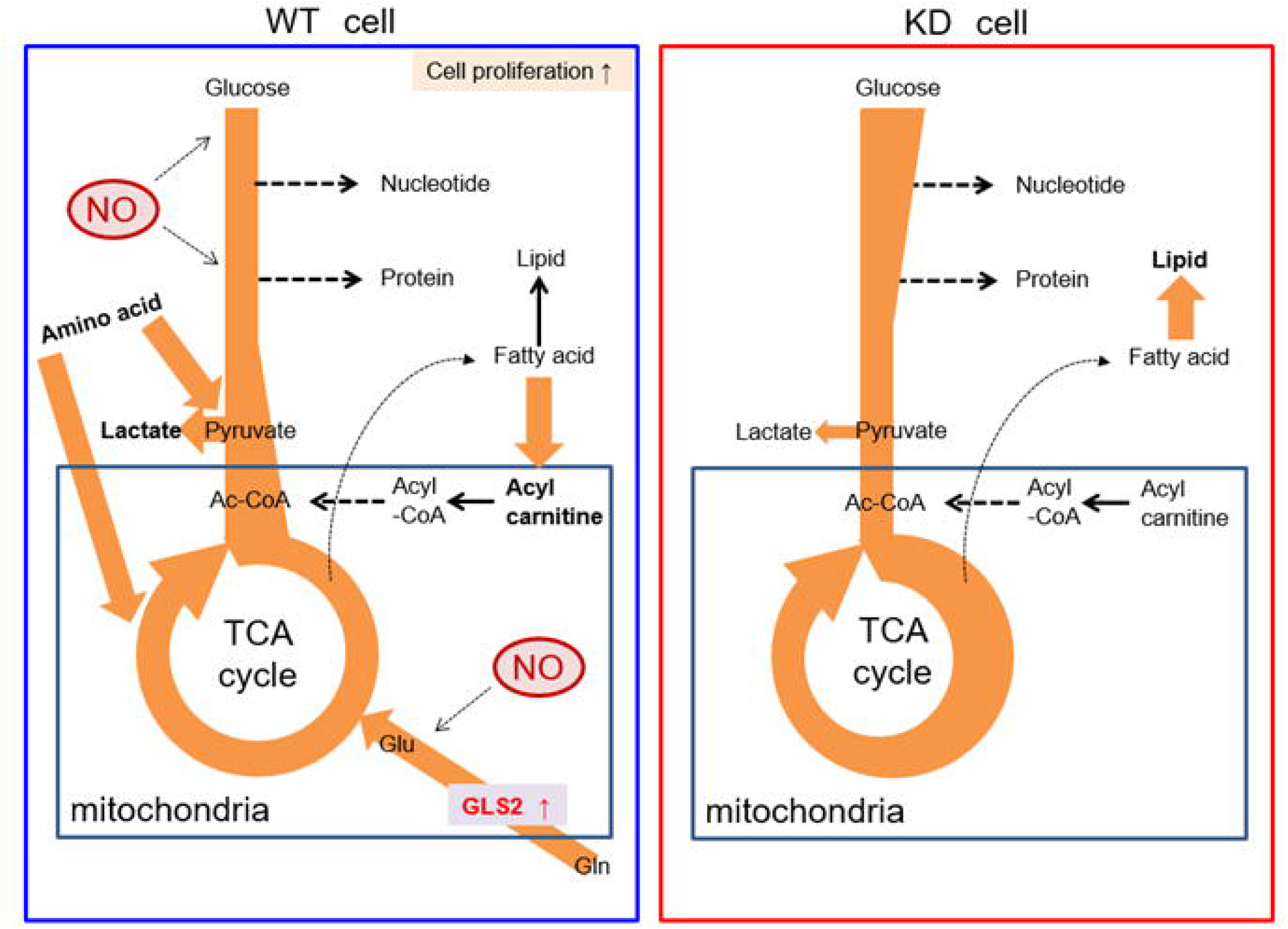
Schematic model of metabolic flux in SW480 cells and TMEM180 KD derivatives Metabolic fluxes revealed in this study are shown. Arrow direction indicates metabolite flux. Bold type indicates a large quantity of metabolites. Nitric oxides are shown as red circle, and glutaminase GLS2 is shown in red font.

## Acknowledgements

The authors thank members of the Matsumura laboratory for helpful discussion and H. Shindo and M. Shimada for their secretarial support. The authors also thank Dr. Y. Uda (Human Metabolome Technologies, Inc.) for analytical support of metabolomics data. This work was financially supported in part by a Research and Development grant by the New Energy and Industrial Technology Development Organization (NEDO) to Y.M.; the National Cancer Center Research and Development Fund (26-A-14, 29-A-9, 29-S-1 to Y.M. and 26-A-12 to M.Y.; a Project for Cancer Research and Therapeutic Evolution from the Japan Agency for Medical Research and Development (AMED) (17cm0106415h0002) to Y.M.; and JSPS KAKENHI (JP19K16730) from the Ministry of Education, Culture, Sports, Science and Technology of Japan to T.A.

## Author Contributions

Y.M. provided the original concept for the study. T.A. and Y.M. designed the experiments. T.A. performed most of the experiments with assistance from S.S. and analyzed the data. Y.O. and H.K. performed the experiments for the cell respiration rate assay. M.Y. contributed initial studies of metabolomics. T.A. and Y.M. discussed and wrote the manuscript. All authors reviewed and contributed to the final version of the manuscript.

