## Supplemental Data for "TMEM180 contributes to colorectal cancer proliferation through intracellular metabolic pathways"

**Title:**

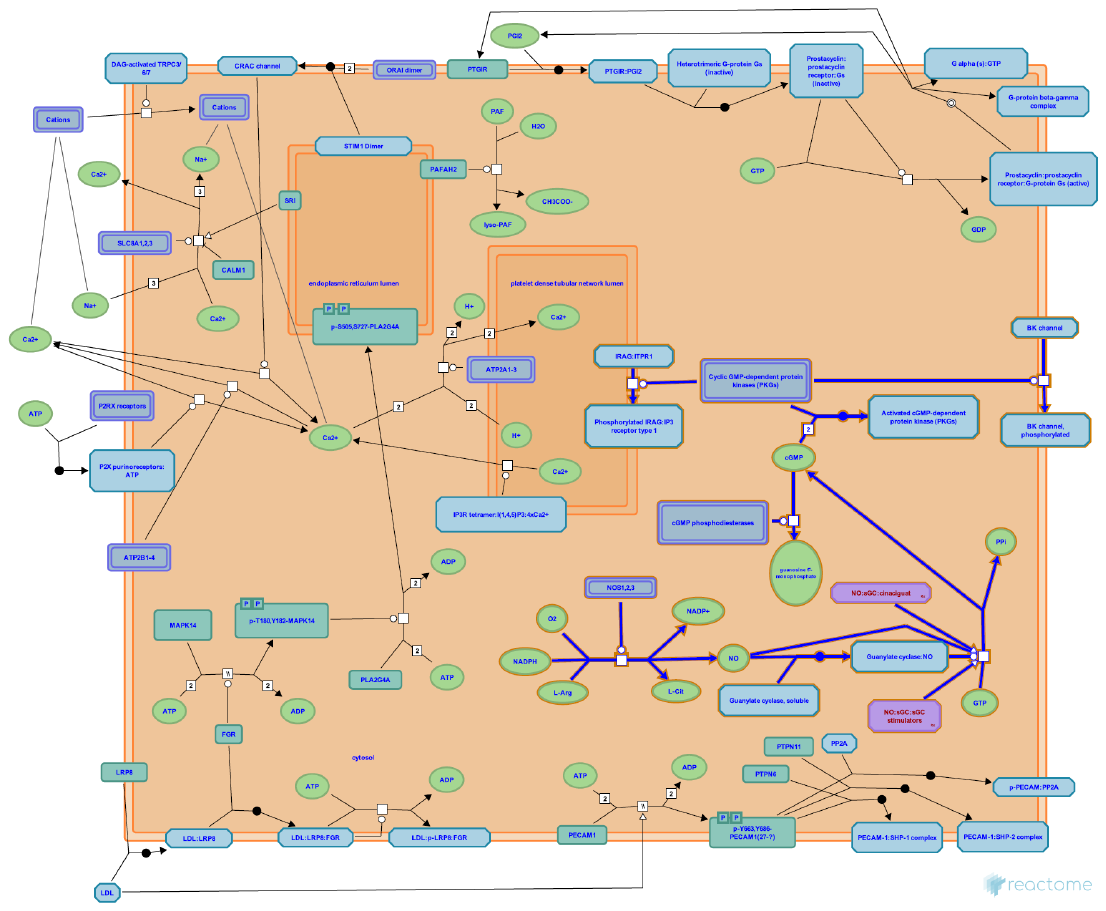

Fig. S1. Nitric oxide stimulates guanylate cyclase

Nitric oxide stimulates guanylate cyclase, from the Reactome pathway map (R-HAS-392154).

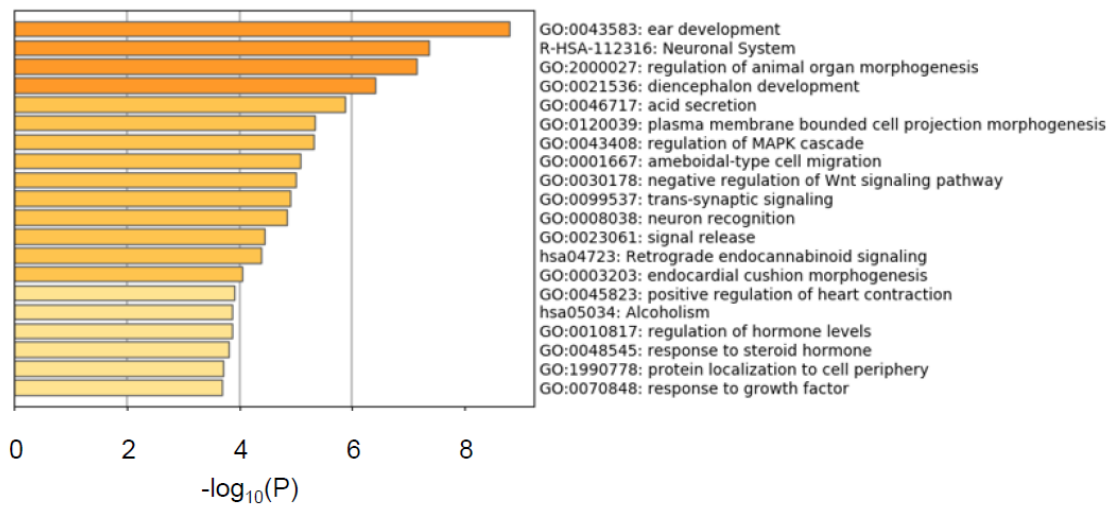

Fig. S2. Enrichment analysis of RNA-seq data

Function enrichment analysis plot, obtained from Metascape, of DEGs inversely correlated with TMEM180 KD from between WT and KD cell samples was shown.

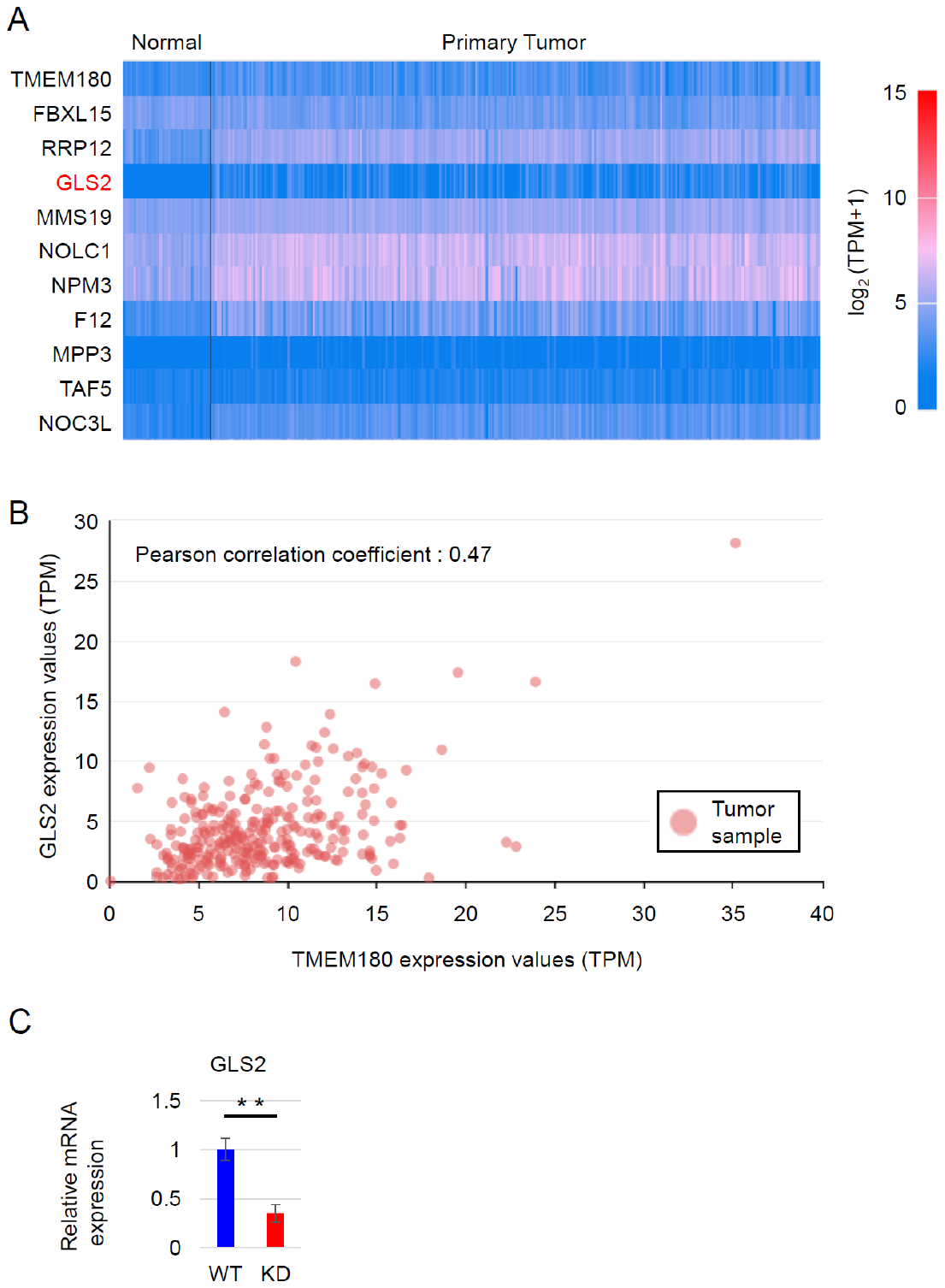

Fig. S3. TCGA data analysis of genes correlated with TMEM180 expression

A.	Heatmap showing the top 10 genes most positively correlated with TMEM180 in TCGA colon adenocarcinoma dataset. Expression level of genes is represented as log_2_(TPM+1).

B.	Gene expression correlation mapping between GLS2 (Y-axis) and TMEM180 (X-axis) in tumor samples in the TCGA colon adenocarcinoma dataset.

C.	Relative expression of GLS2 from our RNA-seq data. WT value is defined as 1. **P < 0.01. Bars = SD.

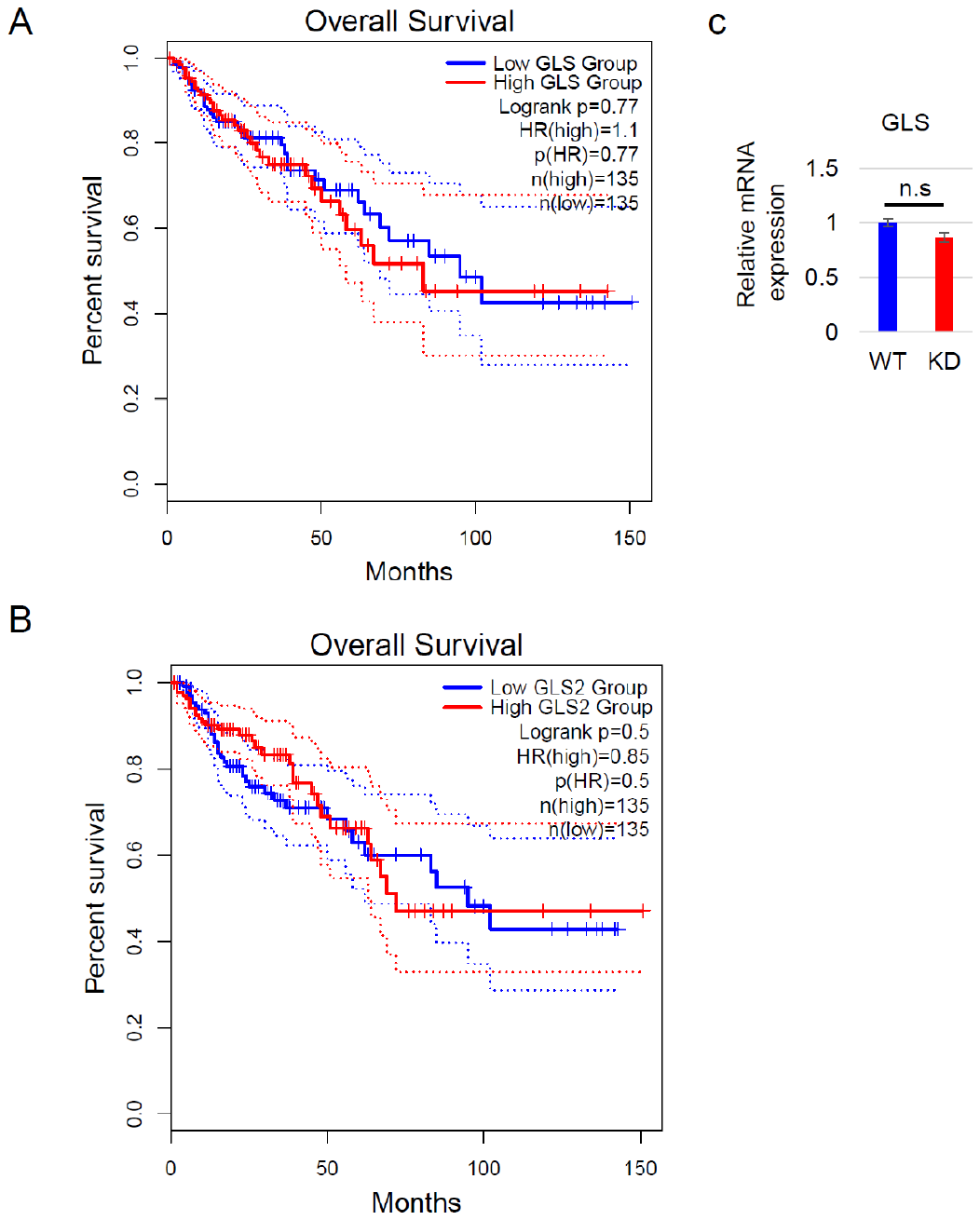

Fig. S4. TCGA analysis of genes correlated with TMEM180 expression

A,B.	Kaplan–Meier plot of patient overall survival stratified by GLS (a) and GLS2 (b) expression. The high-expression group (n=135) is shown in red, and the low-expression group (n=135) is shown in blue.

C.	Relative expression of GLS from our RNA-seq data. WT value is defined as 1. n.s. = not significant. Bars = SD.

Supplementary Table S1. Metabolites in HCA analysis

| Metabolites in cluster A | Metabolites in cluster B |
| --- | --- |
| AC(14:1)-1 | Lactosylceramide(d18:1/24:1)-2 |
| LPS(18:0) | 6-Phosphogluconic acid |
| AC(16:1) | Glucose 6-phosphate |
| AC(18:1) | 1-Stearoyl-glycero-3-phosphocholine-2 |
| Palmitoylcarnitine | 1,2-Dipalmitoyl-glycero-3-phosphoglycerol |
| Thr | NADPH |
| Ser | Galactose 1-phosphate |
| Gln | Betaine aldehyde |
| β-Ala | *cis-Aconitic acid* |
| Gly | 1-Stearoyl-glycero-3-phosphocholine-1 |
| γ-Aminobutyric acid | Isocitric acid |
| Hydroxyproline | Citric acid |
| Ala | Glycerol 3-phosphate |
| Cystathionine | 1-Palmitoyl-glycero-3-phosphoethanolamine LPE(16:0) |
| Asn | 2,3-Diphosphoglyceric acid |
| Asp | Fructose 6-phosphate |
| Glutathione (GSH) | Sedoheptulose 7-phosphate |
| Creatinine | Phosphocreatine |
| Lactosylceramide(d18:1/24:1)-1 | Fructose 1-phosphate |
| Sphinganine | HMG CoA |
| AC(12:0) | Ribose 1-phosphate |
| *N-Acetylglutamic acid* | Folic acid |
| AC(14:0) | 1-Myristoyl-glycero-3-phosphocholine-1 |
| Glu | Fructose 1,6-diphosphate |
| AC(12:1) | 1-Palmitoyl-glycero-3-phosphocholine-1 |
| AC(10:0) | 1-Palmitoyl-glycero-3-phosphocholine-2 |
| His | Betaine |
| Val | cAMP |
| Ile | Ricinoleic acid |
| Met | 1-Oleoyl-glycero-3-phosphocholine-2 |
| Trp | Citrulline |
| Leu | LPE(18:1)-2 |
| Tyr | UDP-glucose |
| Oleic acid | NADH |
| Carnosine | LPE(18:1)-1 |
| Phe | Succinic acid |
|  | 2-Hydroxyglutaric acid |
|  | 2-Phosphoglyceric acid |
|  | Glyceraldehyde 3-phosphate |
|  | NAD^+^ |
|  | Xylulose 5-phosphate |
|  | ADP-ribose |
|  | Phosphoenolpyruvic acid |
|  | Dihydroxyacetone phosphate |
|  | 3-Phosphoglyceric acid |
|  | 1-Myristoyl-glycero-3-phosphocholine-2 |
|  | IMP |
|  | Adenine |

Supplementary Table S2. Metabolites in PCA analysis with positive values in PC1

| Rank | Metabolites | R | p-value |
| --- | --- | --- | --- |
| 1 | 1-Palmitoyl-glycero-3-phosphocholine-2 | 0.982 | 4.58E-04 |
| 2 | Fructose 1,6-diphosphate | 0.963 | 1.99E-03 |
| 3 | Ricinoleic acid | 0.960 | 2.34E-03 |
| 4 | Betaine | 0.959 | 2.49E-03 |
| 5 | Glycerol 3-phosphate | 0.954 | 3.13E-03 |
| 6 | cAMP | 0.950 | 3.66E-03 |
| 7 | 6-Phosphogluconic acid | 0.946 | 4.23E-03 |
| 8 | 1-Myristoyl-glycero-3-phosphocholine-1 | 0.946 | 4.24E-03 |
| 9 | Citrulline | 0.938 | 5.67E-03 |
| 10 | 1-Palmitoyl-glycero-3-phosphocholine-1 | 0.921 | 9.19E-03 |
| 11 | Isocitric acid | 0.920 | 9.31E-03 |
| 12 | 1-Stearoyl-glycero-3-phosphocholine-1 | 0.915 | 1.04E-02 |
| 13 | Lactosylceramide(d18:1/24:1)-2 | 0.910 | 1.19E-02 |
| 14 | 1-Stearoyl-glycero-3-phosphocholine-2 | 0.908 | 1.22E-02 |
| 15 | 1,2-Dipalmitoyl-glycero-3-phosphoglycerol | 0.894 | 1.64E-02 |
| 16 | 1-Myristoyl-glycero-3-phosphocholine-2 | 0.889 | 1.78E-02 |
| 17 | Glucose 6-phosphate | 0.885 | 1.92E-02 |
| 18 | Citric acid | 0.885 | 1.92E-02 |
| 19 | Betaine aldehyde | 0.869 | 2.45E-02 |
| 20 | LPE(18:1)-1 | 0.865 | 2.60E-02 |
| 21 | 1-Oleoyl-glycero-3-phosphocholine-2 | 0.864 | 2.64E-02 |
| 22 | *cis-Aconitic acid* | 0.858 | 2.89E-02 |
| 23 | Succinic acid | 0.842 | 3.53E-02 |
| 24 | Galactose 1-phosphate | 0.816 | 4.74E-02 |
| 25 | Fructose 6-phosphate | 0.810 | 5.09E-02 |
| 26 | UDP-glucose | 0.808 | 5.18E-02 |
| 27 | 2-Hydroxyglutaric acid | 0.785 | 6.42E-02 |
| 28 | NAD^+^ | 0.783 | 6.54E-02 |
| 29 | HMG CoA | 0.781 | 6.68E-02 |
| 30 | Glyceraldehyde 3-phosphate | 0.778 | 6.86E-02 |

Supplementary Table S3. Metabolites in PCA analysis with negative values in PC1

| Rank | Metabolites | R | p-value |
| --- | --- | --- | --- |
| 30 | Ile | -0.895 | 1.60E-02 |
| 29 | Leu | -0.900 | 1.46E-02 |
| 28 | Trp | -0.900 | 1.45E-02 |
| 27 | AC(14:0) | -0.900 | 1.44E-02 |
| 26 | His | -0.901 | 1.43E-02 |
| 25 | AC(10:0) | -0.911 | 1.17E-02 |
| 24 | Glu | -0.916 | 1.04E-02 |
| 23 | LPS(18:0) | -0.922 | 8.99E-03 |
| 22 | Putrescine | -0.925 | 8.30E-03 |
| 21 | AC(12:0) | -0.926 | 7.96E-03 |
| 20 | Sphingosine | -0.929 | 7.34E-03 |
| 19 | AC(14:1)-1 | -0.930 | 7.26E-03 |
| 18 | Met | -0.936 | 6.08E-03 |
| 17 | Lactosylceramide(d18:1/24:1)-1 | -0.936 | 5.92E-03 |
| 16 | Asp | -0.937 | 5.83E-03 |
| 15 | Creatinine | -0.942 | 4.98E-03 |
| 14 | Glutathione (GSH) | -0.943 | 4.84E-03 |
| 13 | Sphinganine | -0.967 | 1.66E-03 |
| 12 | AC(18:1) | -0.971 | 1.25E-03 |
| 11 | Hydroxyproline | -0.979 | 6.54E-04 |
| 10 | Palmitoylcarnitine | -0.980 | 6.13E-04 |
| 9 | γ-Aminobutyric acid | -0.982 | 4.70E-04 |
| 8 | Ala | -0.987 | 2.47E-04 |
| 7 | Asn | -0.988 | 2.18E-04 |
| 6 | Gln | -0.991 | 1.10E-04 |
| 5 | β-Ala | -0.992 | 8.54E-05 |
| 4 | Cystathionine | -0.994 | 5.14E-05 |
| 3 | Thr | -0.996 | 1.89E-05 |
| 2 | Gly | -0.997 | 1.39E-05 |
| 1 | Ser | -0.999 | 1.75E-06 |
